# Amyloid, Tau, and APOE in Alzheimer’s Disease: Impact on White Matter Tracts

**DOI:** 10.1101/2024.08.05.606560

**Authors:** Bramsh Qamar Chandio, Julio E. Villalon-Reina, Talia M. Nir, Sophia I. Thomopoulos, Yixue Feng, Sebastian Benavidez, Neda Jahanshad, Jaroslaw Harezlak, Eleftherios Garyfallidis, Paul M. Thompson, the Alzheimer’s Disease Neuroimaging Initiative

## Abstract

Alzheimer’s disease (AD) is characterized by cognitive decline and memory loss due to the abnormal accumulation of amyloid-beta (A*β*) plaques and tau tangles in the brain; its onset and progression also depend on genetic factors such as the apolipoprotein E (APOE) genotype. Understanding how these factors affect the brain’s neural pathways is important for early diagnostics and interventions. Tractometry is an advanced technique for 3D quantitative assessment of white matter tracts, localizing microstructural abnormalities in diseased populations *in vivo*. In this work, we applied BUAN (Bundle Analytics) tractometry to 3D diffusion MRI data from 730 participants in ADNI3 (phase 3 of the Alzheimer’s Disease Neuroimaging Initiative; age range: 55-95 years, 349M/381F, 214 with mild cognitive impairment, 69 with AD, and 447 cognitively healthy controls). Using along-tract statistical analysis, we assessed the localized impact of amyloid, tau, and APOE genetic variants on the brain’s neural pathways. BUAN quantifies microstructural properties of white matter tracts, supporting along-tract statistical analyses that identify factors associated with brain microstructure. We visualize the 3D profile of white matter tract associations with tau and amyloid burden in Alzheimer’s disease; strong associations near the cortex may support models of disease propagation along neural pathways. Relative to the neutral genotype, APOE *ɛ*3/*ɛ*3, carriers of the AD-risk conferring APOE *ɛ*4 genotype show microstructural abnormalities, while carriers of the protective *ɛ*2 genotype also show subtle differences. Of all the microstructural metrics, mean diffusivity (MD) generally shows the strongest associations with AD pathology, followed by axial diffusivity (AxD) and radial diffusivity (RD), while fractional anisotropy (FA) is typically the least sensitive metric. Along-tract microstructural metrics are sensitive to tau and amyloid accumulation, showing the potential of diffusion MRI to track AD pathology and map its impact on neural pathways.

## 1. Introduction

Alzheimer’s disease (AD) is a neurodegenerative disorder characterized by progressive cognitive decline and memory loss. Central to its pathology are the abnormal accumulation of amyloid-beta (*Aβ*) plaques and tau tangles in the brain.^1–3^ The onset and progression of these pathological processes are influenced by genetic factors such as the apolipoprotein E (APOE) gene.^4^ AD pathology not only affects gray matter but also profoundly impacts white matter tracts, which serve as the brain’s communication mechanism; these tracts connect different brain regions and facilitate efficient signal transmission. Understanding how amyloid, tau, and APOE influence white matter integrity is crucial for developing early diagnostic tools and monitoring the effects of targeted interventions on the brain.

Amyloid-beta peptides aggregate to form plaques, primarily affecting gray matter^5^ but also extending to white matter tracts by disturbing cellular function.^6,7^ *Aβ* deposition leads to myelin degradation, which disrupts the insulating layer around nerve fibers, and axonal injury, which impairs neurons’ ability to communicate effectively. The presence of *Aβ* can trigger a chronic inflammatory response, worsening white matter damage through microglial activation and the release of pro-inflammatory cytokines.

Tau is a microtubule-associated protein that stabilizes microtubules in neurons. In AD, tau becomes hyperphosphorylated and forms neurofibrillary tangles,^8^ affecting microtubule stability, disrupting axonal transport, and impairing neuronal function. The propagation of tau pathology correlates with synaptic loss and neuronal degeneration, affecting both gray and white matter regions and leading to widespread brain dysfunction.^9,10^

The apolipoprotein E (APOE) gene has three common variants: *ɛ*2, *ɛ*3, and *ɛ*4. APOE *ɛ*2 is the least common, and carriers have a lower risk of developing AD. It may have a protective effect on white matter structure,^11,12^ leading to less degeneration compared to those with APOE *ɛ*3 or *ɛ*4, possibly due to enhanced lipid metabolism and repair mechanisms. APOE *ɛ*3 is the most common variant and is considered neutral, while the APOE *ɛ*4 variant is the greatest known common genetic risk factor for late-onset AD, roughly tripling lifetime risk of AD per allele carried.^13^ APOE *ɛ*4 is less effective in clearing *Aβ* from the brain, leading to greater *Aβ* plaque accumulation and subsequent white matter damage.^14^ APOE is involved in lipid transport and metabolism, essential for myelin maintenance. The *ɛ*4 variant affects these processes, leading to compromised myelin maintenance. APOE *ɛ*4 carriers also exhibit increased inflammation and vascular dysfunction, contributing to white matter lesions and impaired cerebral blood flow. Even so, relatively little is known about the 3D profile of APOE effects on white matter microstructure.

Investigating the impact of amyloid, tau, and APOE on white matter is vital for advancing AD research and treatment. White matter metrics may deteriorate before gray matter atrophy and clinical symptoms appear, making them a potential biomarker for early detection or monitoring interventions. Diffusion MRI, being less invasive than PET, could offer a better alternative for tracking disease progression if metrics sensitive to these pathologies are identified. Identifying white matter tracts affected may also help to evaluate targeted therapies to protect or restore these pathways. Furthermore, understanding how different APOE genotypes affect white matter may lead to personalized therapeutic strategies, improving outcomes for individuals with specific genetic profiles.

Diffusion MRI^15–17^ measures water diffusion in the brain, revealing the microstructural properties of the underlying tissue. Tractography, derived from diffusion MRI data,^18–20^ maps and visualizes white matter pathways by tracking the directional profiles of water diffusion, providing a detailed picture of brain connectivity. Tractometry enhances this by quantifying specific microstructural properties, such as fractional anisotropy (FA) or mean diffusivity (MD), along the length of individual tracts. This technique maps microstructural alterations in the brain’s white matter tracts.^21–25^ It analyzes the coherence of neural connections, allowing for precise assessment of characteristic changes in neurological conditions such as Alzheimer’s disease or Parkinson’s disease.^26^

White matter (WM) microstructure changes with age, and there is a regional variation in the age-dependent trajectories of maturation and decline for the major white matter metrics across the lifespan.^27,28^ Several studies of regional microstructure in Alzheimer’s disease have used tract-based spatial statistics (TBSS),^29^ to link microstructural metrics in specific brain regions to amyloid positivity and clinical dementia severity.^30–32^ However, the resolution of TBSS maps is limited by the regions defined in the atlases used.^29^ To address this, tractometry methods such as BUAN (Bundle Analytics)^33^ map microstructural parameters along the length of white matter tracts, mapping disease effects on neural pathways in 3D and at a finer anatomical scale.^23,25,26,34,35^ Recently, Ba Gari *et al.*^34^ used a tractography-based medial tract analysis (MeTa) to enhance the sensitivity for detecting associations of AD, amyloid and tau with DTI microstructural metrics, compared to TBSS.

In this study, we applied our advanced tractometry method, BUAN (Bundle Analytics), to evaluate the impact of amyloid, tau, APOE *ɛ*4, and APOE *ɛ*2 on the microstructure of the brain’s white matter tracts. BUAN maps the microstructural properties of white matter tracts, and fits along-tract statistical models to detect effects on microstructure that are associated with amyloid plaques, tau tangles, and different APOE genotypes. This is crucial for understanding the effects of AD pathology on brain connectivity. Overall, we found that a range of microstructural metrics were sensitive to tau and amyloid, the two key biomarkers for detecting Alzheimer’s disease, supporting the role of diffusion MRI as a non-invasive measure of AD pathology. Relative to APOE *ɛ*3/*ɛ*3 carriers, microstructural alterations were also identified in APOE *ɛ*4 carriers and to a lesser extent in *ɛ*2 carriers.

Overall, mean diffusivity (MD) was most strongly associated with AD pathology, followed by axial diffusivity (AxD) and radial diffusivity (RD). Fractional anisotropy (FA) was the least sensitive metric. The tendency to detect stronger associations in tract regions closer to the cortex may support propagative or “epidemic spreading” models of AD pathology,^36^ which argue that AD pathology spreads dynamically along neural pathways or in functionally synchronous networks; future longitudinal studies are needed to verify this.

## 2. Methods

### 2.1. Data

Data from 730 ADNI3 participants (phase 3 of the Alzheimer’s Disease Neuroimaging Initiative; age range: 55-95 years, 349M/381F, 214 with mild cognitive impairment (MCI), 69 with AD, and 447 cognitively healthy controls (CN)) scanned with 7 acquisition protocols (GE36, GE54, P33, P36, S127, S31, S55) were included. Tables 1 and 2 in Fig. 2 detail demographic and acquisition protocol information. A*β*-status, i.e., positive (A*β*+) or negative (A*β*–), was determined by either mean 18F-florbetapir (A*β*+ defined as *>*1.11)^37,38^ or florbetaben (A*β*+ defined as *>*1.20)^39,40^ PET cortical SUVR uptake, normalized by using a whole cerebellum reference region. Tau positivity was defined as a tau SUVR *>* 1.23.

**Fig. 1.**
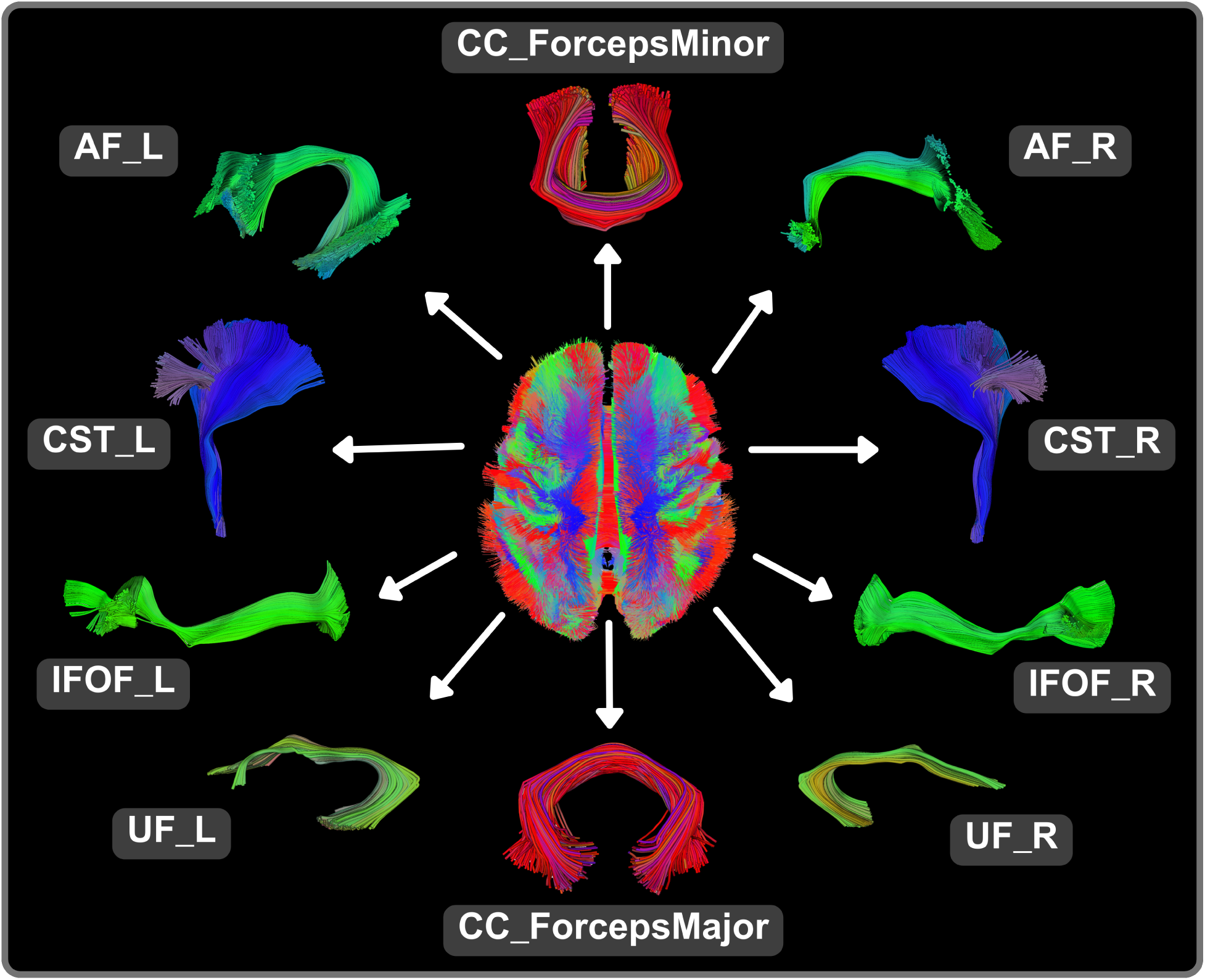
The brain contains millions of axonal connections; the trajectories of the major neural pathways can be digitally reconstructed into a whole brain tractogram using diffusion MRI and tractography techniques. Specific white matter tracts are extracted for visualization and detailed analysis, offering a more localized and focused examination of pathways within the brain.

**Fig. 2.**
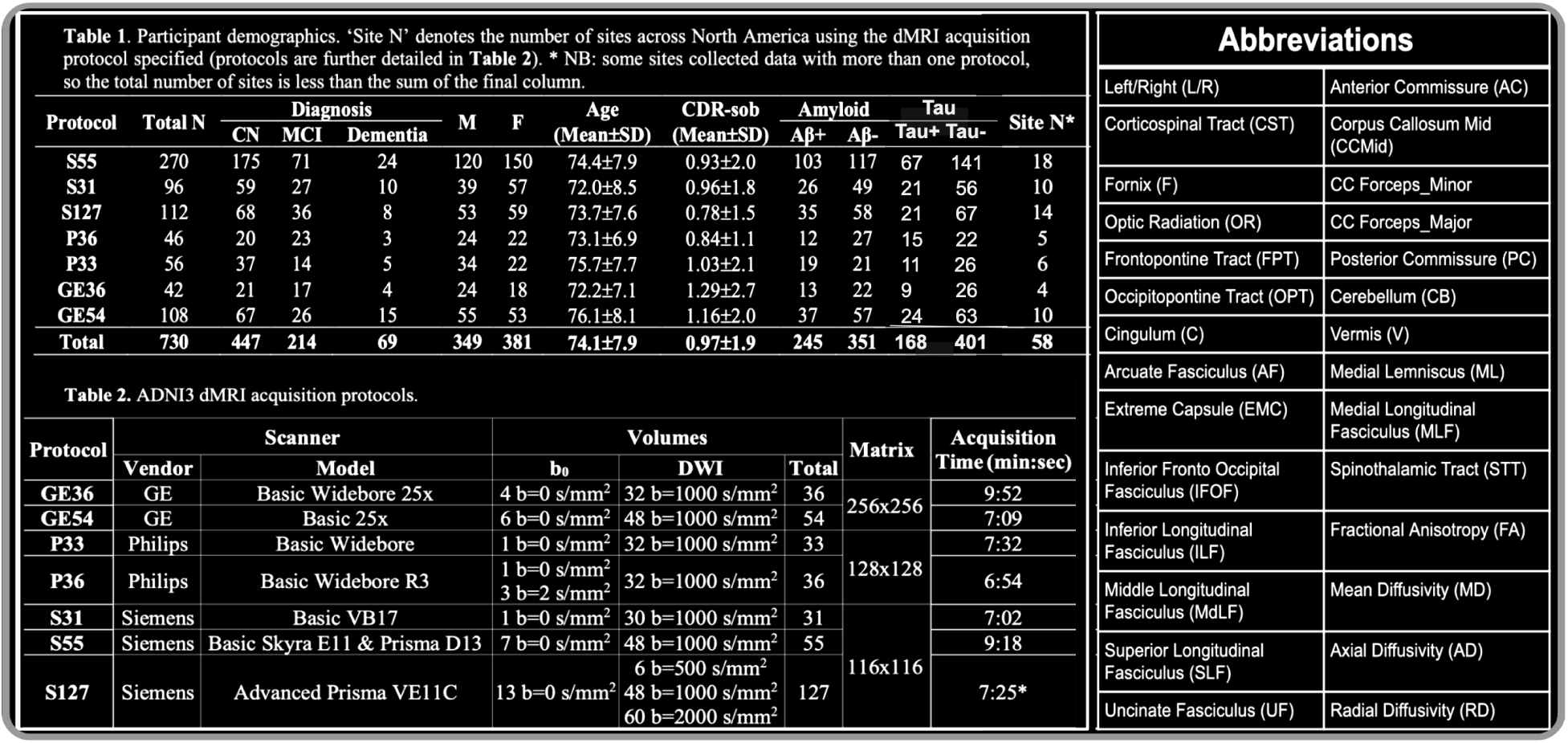
Tables 1 and 2 detail demographic and scanner protocol information for the ADNI3 data used in our experiments (data from Thomopoulos et al, 2021). The abbreviation table on the right lists the 38 white matter tracts and four microstructural measures analyzed in this work.

### 2.2. Diffusion MRI Processing

Raw diffusion MRI (dMRI) were preprocessed using the ADNI3 dMRI protocol.^41,42^ Preprocessing of raw diffusion MRI (dMRI) data involved several steps: denoising raw dMRI data using DIPY’s principal component analysis (PCA) for GE data, and Marchenko-Pastur PCA for Siemens and Philips data denoising.^43,44^ Gibbs artifacts were corrected using MRtrix’s *degibbs* tool,^45,46^ and extracerebral tissue was removed (skull stripping) with FSL’s BET.^47,48^ Eddy currents and motion were corrected using FSL’s eddy cuda tool with additional corrections for slice-to-volume and outlier detection.^48,49^ Bias field inhomogeneities were corrected using MRtrix’s *dwibiascorrection* ANTS function. Preprocessed T1w images from the ADNI database were further processed and aligned to the dMRI data.^46,50^ ADNI3 dMRI data lacked reversed phase-encode blips, so echo-planar imaging (EPI) distortion corrections were made using nonlinear registrations to T1-weighted anatomical images. The processed dMRI data were converted back to native space through a series of inversions of the registration matrices, with final outputs visually inspected and manually adjusted as necessary. The diffusion tensor imaging (DTI) model was used to extract 4 microstructural measures from processed dMRI: fractional anisotropy (FA), mean, axial, and radial diffusivity (MD, AxD, RD).

### 2.3. BUAN Tractometry

We applied a robust and unbiased model-based spherical deconvolution^51^ reconstruction method and a probabilistic particle filtering tracking algorithm that uses tissue partial volume estimation (PVE) to reconstruct ^52^ whole-brain tractograms. For tracking, the seed mask was created from the white matter (WM) PVE (WM PVE *>* 0.5), seed density per voxel was set to 2, and step size was set to 0.5. We extracted 38 white matter (WM) tracts from tractograms using auto-calibrated RecoBundles ^23,53^ (see Fig. 2 for full names) using model bundles from the HCP-842 tractography atlas.^54^

After extracting WM bundles, we nonlinearly registered each subject’s bundles to model bundles in MNI-space using a streamline-based nonlinear registration method, BundleWarp.^55^ Optimal registration of tracts to atlas bundles is crucial for finding accurate segment correspondences among subjects and populations. This enhances the sensitivity of group statistical analyses by eliminating errors due to misalignment across subjects.

BUAN creates the bundle profiles for each bundle using 4 DTI-based microstructural metrics: FA, MD, RD, and AxD calculated in the diffusion native space (see Figure 2 for full bundle names). Bundle profiles are created by dividing the bundles into 100 horizontal segments using the model bundle centroids along the length of the tracts in common space. We cluster our model bundles using the QuickBundles^56^ method to obtain a cluster centroid consisting of 100 points per centroid. We calculate Euclidean distances between every point on every streamline of the bundle and 100 points in the model bundle centroid. A segment number is assigned to each point in a bundle based on the shortest distance to the nearest model centroid point. The streamlines are not resampled to have a specific number of points, and we do not change the distribution of points. Since the assignment of segment numbers is performed in the common space, we establish the segment correspondence among subjects from different groups and populations. Microstructural measures such as FA are then projected onto the points of the bundles in native space. Note that the nonlinearly moved bundles are only used to assign segment numbers to streamlines (and points on the streamlines) in the bundles. Actual statistical analysis always takes place in the native space of the diffusion data. The statistical analysis step uses bundles of the original shape and microstructural measures in the native space using segment labels given during the assignment step for segment-specific group analysis.

Bundle profiles are harmonized using the ComBat method^57,58^ to correct for scanner/site effects as described in the harmonized BUAN tractometry pipeline.^59^ After data harmonization, we assume each bundle type has its own data distribution, which is considered independent of the rest of the bundles in the brain. For each tract and metric, we pool bundle profiles for a given tract across all subjects from CN, MCI, and AD groups. Pooled bundle profiles consist of 100 segments, and each segment is modeled as a feature. Linear Mixed Models are applied to WM bundles; age and sex are modeled as fixed effects and scanner and subject as a random effect term, the response variable being each DTI metric. Though we harmonized the profiles with ComBat, we further account for scanner and/or site effects by adding it as a random term in the linear mixed models (LMMs)^60^ to eliminate any remaining artifacts contributed by scanner/site. We used FURY^61^ software to visualize tractometry results in this paper.

### 2.4. Statistics

We used LMMs to test the effects of amyloid positivity, tau positivity, and different APOE variants on 38 white matter tracts. In each experiment, age and sex were modeled as fixed effects, and the scanner and subject were modeled as random terms.

#### 2.4.1. Multiple test corrections

Multiple testing correction is a statistical adjustment process that can control the rate or likelihood of false positives when performing numerous simultaneous tests.^62^ In neuroimaging studies, where thousands of brain regions or voxels are analyzed for significant differences or correlations, this adjustment is crucial. It ensures the integrity and reliability of the results by controlling the overall rate of false positives. Common correction methods include the Bonferroni correction,^63^ which is stringent and adjusts the significance threshold by dividing it by the number of tests, and the False Discovery Rate (FDR)^64^ method, which limits the proportion of false positives among significant findings. These corrections ensure that detected effects are truly significant and not due to random variation.

As white matter tracts generated by tractography are not as extensively studied as voxel or ROI-based methods, selecting the appropriate multiple testing correction is challenging. We divided each bundle into 100 segments; for tract-specific FDR correction, we use 100 *p*-values per bundle to correct for multiple tests using the FDR method. We refer to this bundle-specific FDR corrected threshold as the local threshold, as it only depends on statistics within that bundle. Additionally, we performed multiple test corrections across all bundles in the brain by pooling 100 *p*values from each of the 38 tracts, yielding a total of 3,800 *p*values to determine the global FDR-corrected threshold. We consider tract effects to be significant if they pass both local and global FDR thresholds.

## 3. Results

We ran the following five experiments to detect associations of various variables on 38 white matter tracts of the brain. We tested microstructural associations (1) with amyloid positivity; (2) with tau positivity; (3) comparing non *ɛ*4 carriers *ɛ*2*ɛ*3/*ɛ*3*ɛ*3/*ɛ*2*ɛ*2 with subjects carrying at least one *ɛ*4 gene; *ɛ*2*ɛ*4/*ɛ*3*ɛ*4/*ɛ*4*ɛ*4, (4) comparing *ɛ*3*ɛ*3 with *ɛ*3*ɛ*4/*ɛ*4*ɛ*4, and (5) comparing *ɛ*3*ɛ*3 with *ɛ*3*ɛ*2/*ɛ*2*ɛ*2.

As an overview of the results, quantitative quantile-quantile (QQ) plots (Fig. 3) summarize the overall association signal detected across all 38 white matter bundles between each of the biomarkers (amyloid, tau, and APOE) and each of the DTI metrics (FA, MD, RD, and AxD). These plots visually represent the strength of associations between these biomarkers and DTI metrics, helping to identify which combinations show the most significant relationships.

**Fig. 3.**
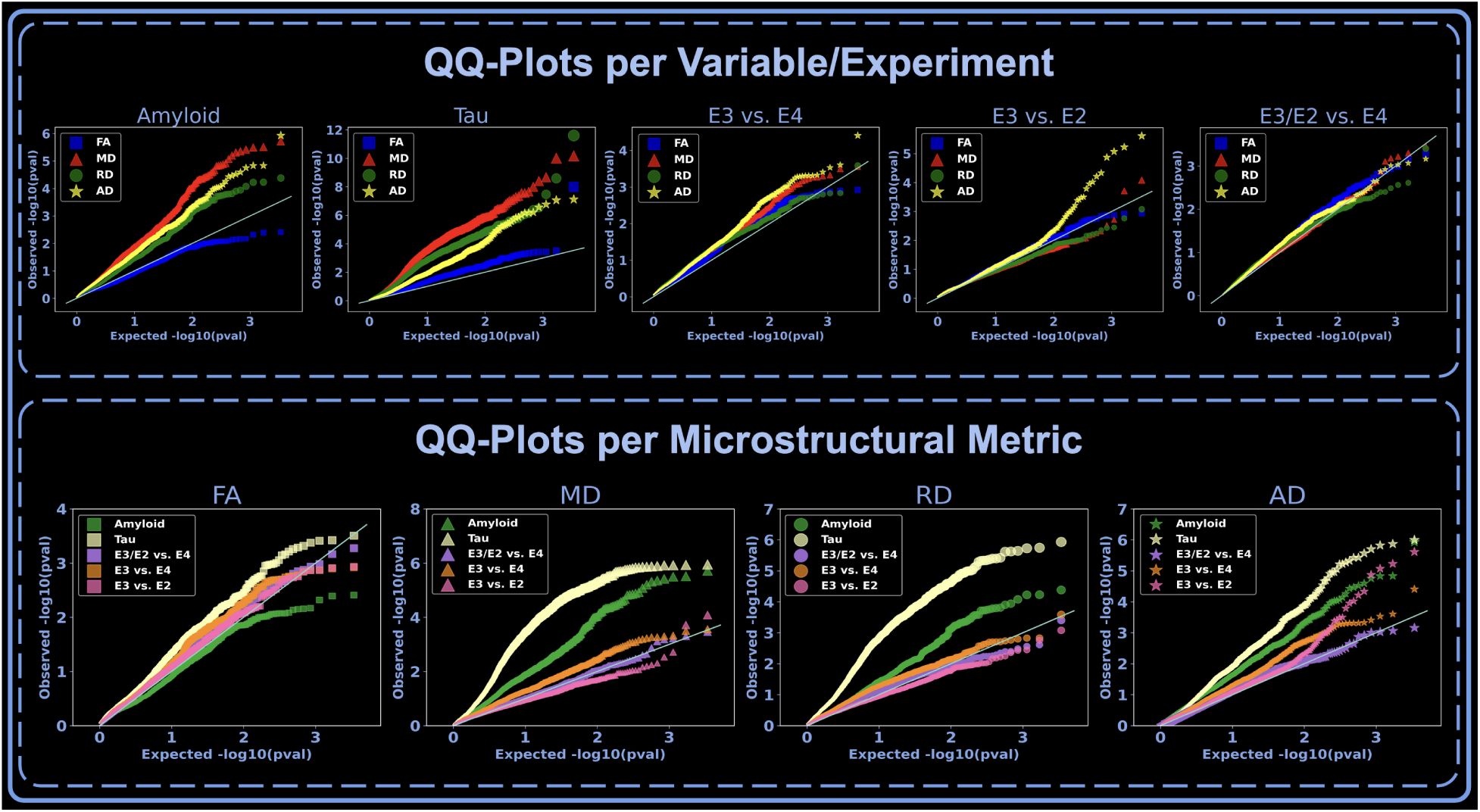
QQ plots summarize the signal detected by each biomarker (amyloid, tau, and ApoE) and DTI metric (FA, MD, RD, and AxD) across all 38 bundles, indicating which biomarkers and metrics show the strongest associations. In the first row, the plots show which metric shows the strongest association for each biomarker. *P*-values of the 38 tracts were pooled together for each DTI metric and visualized in QQ plots. In the second row, we analyze for each metric which biomarker shows significant associations. Note the *y*-axis range varies across the plots depending on the observed range of *p*-values.

In the visualization layout, the first row of QQ plots highlights which DTI metric exhibits the strongest association with each biomarker. Here, the *p*-values of the 38 tracts were pooled for each DTI metric and presented in these plots, allowing for a comprehensive assessment of each metric’s sensitivity to changes in biomarker levels.

The second focuses on the relationship from the opposite perspective: for each DTI metric, it shows which biomarker shows significant associations (the scale of the *y*-axis varies across the QQ plots to adapt to the observed range of *p*-values).

### 3.1. Amyloid

We ran BUAN to assess the effect of amyloid positivity on 38 white matter tracts based on data from 329 amyloid-negative (CN: 235, MCI: 86, Dementia: 8) (156M, 173F) and 277 amyloid-positive (CN: 139, MCI: 87, Dementia: 51) (131M, 146F) participants from the ADNI3 dataset.

We found that the following tracts and measures showed significant differences between amyloid negative and amyloid positive: cingulum left (AxD, MD), corpus callosum forceps major and middle sector (MD, RD), extreme capsule left (MD, RD) and right (AxD, MD, RD), frontopontine tract left (AxD, MD) and right (AxD), inferior longitudinal fasciculus right (AxD, MD, RD), middle longitudinal fasciculus left (AxD, MD) and right (AxD, MD, RD), occipito-pontine tract left (MD), optic radiation right (MD), posterior commissure (AxD), and spinothalamic tract left (RD).

In significant tracts, diffusivity metrics increase while fractional anisotropy decreases, in those with higher levels of amyloid pathology (this is in the same direction as the known effects of dementia on these metrics).

### 3.2. Tau

We ran BUAN to assess the effect of tau positivity on 38 white matter tracts based on data from 401 tau-negative (CN: 293, MCI: 95, and Dementia: 13) (192M, 209F) and 168 amyloid-positive (CN: 60, MCI: 68, and Dementia: 40) (75M, 93F) participants in the ADNI3 dataset.

The following tracts and measures showed significant associations between tau positivity and microstructure:

Arcuate fasciculus left (MD, RD), cingulum left and right (MD, RD), corpus callosum - forceps major (MD, RD), forceps minor (FA, MD, RD) and mid (AD, MD, RD), corticospinal Tract left and right (MD, RD), extreme capsule left and right (AxD, MD, RD), frontopontine tract left (MD, RD) and right (FA, AxD, MD, RD), inferior fronto-occipital fasciculus right (RD), inferior longitudinal fasciculus left (MD, RD) and right (AxD, MD, RD), middle longitudinal fasciculus left (AxD, MD, RD) and right (AxD, FA, MD, RD), occipito-pontine tract left (MD, RD) and right (AxD, MD, RD), optic radiation left (RD) and right (AxD, MD, RD), and uncinate fasciculus right (MD, RD).

In significant tracts, most diffusivity metrics increase while fractional anisotropy decreases, in line with the expected direction of microstructural abnormalities previously reported in dementia. However, in some tracts, changes in AxD vary along the length of the tracts.

In Fig. 4, we compare the impact on white matter tracts as influenced by amyloid and tau. Only those tracts that demonstrate significant effects, meeting both local and global false discovery rate (FDR) criteria for amyloid and tau, are included. Significant associations with each biomarker in conjunction with DTI metrics are highlighted in red. We consistently observe stronger associations with tau across various white matter tracts, as illustrated in the QQ-plot at the right end of the figure. For all metrics assessed, tau shows stronger associations compared to amyloid.

**Fig. 4.**
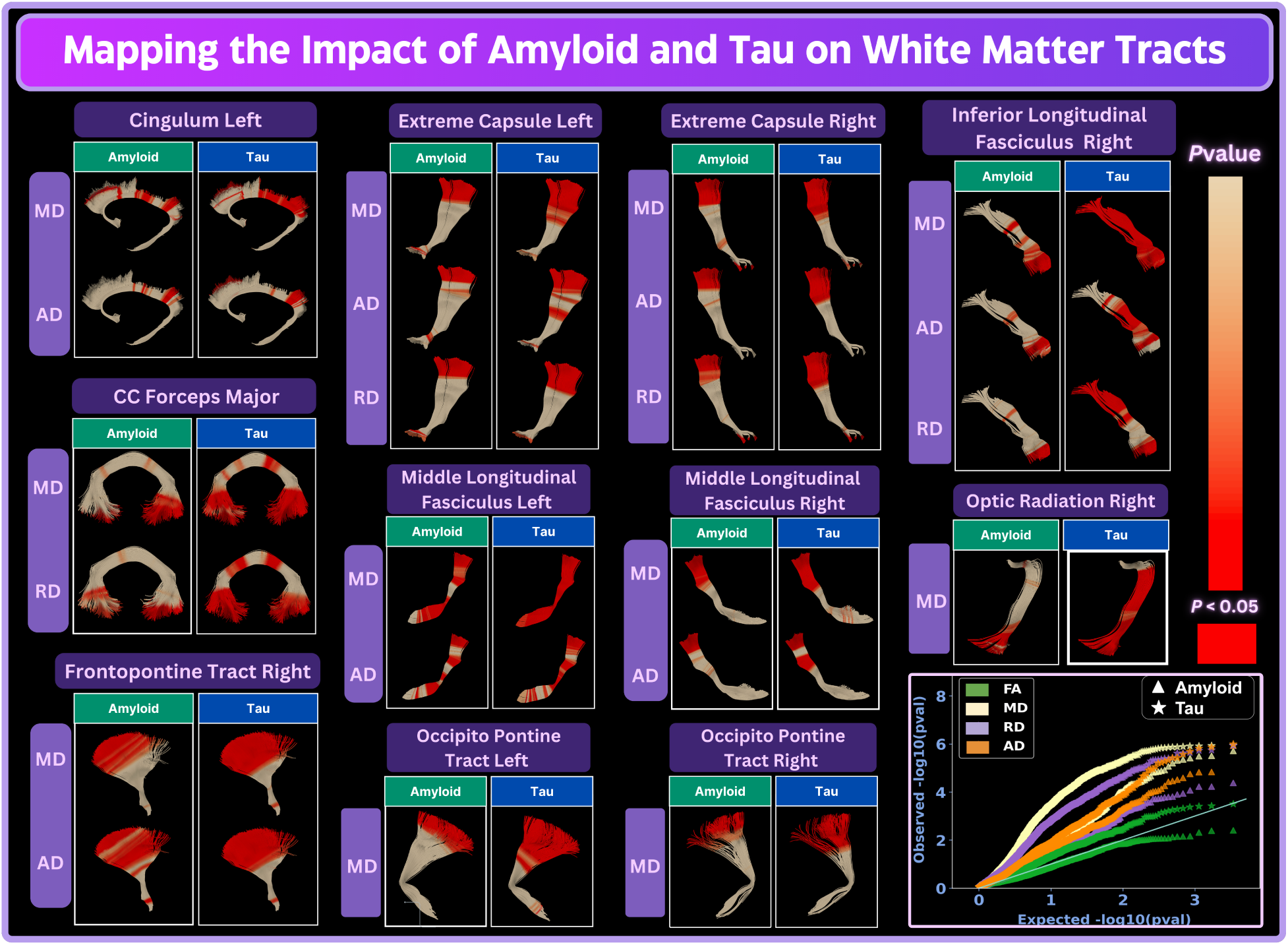
We compare the effects of amyloid and tau on white matter microstructure along the major white matter tracts. Only tracts showing significant effects, passing both local and global FDR for amyloid and tau, are visualized. *Red* highlights significant associations between the measures of Alzheimer’s disease pathology and the microstructural metrics computed with DTI. We consistently observe the strongest associations with tau in various white matter tracts, as seen in the QQ-plot at the right end of the figure. Tau outperforms amyloid in terms of strength of association, for each microstructural metric.

MD metrics exhibit the strongest association signal for both amyloid and tau. We illustrate the localized effects of tau on MD metrics in Fig. 5. Each tract is color-coded based on *p*-values, with tracts showing *p*-values less than 0.05 highlighted in red.

**Fig. 5.**
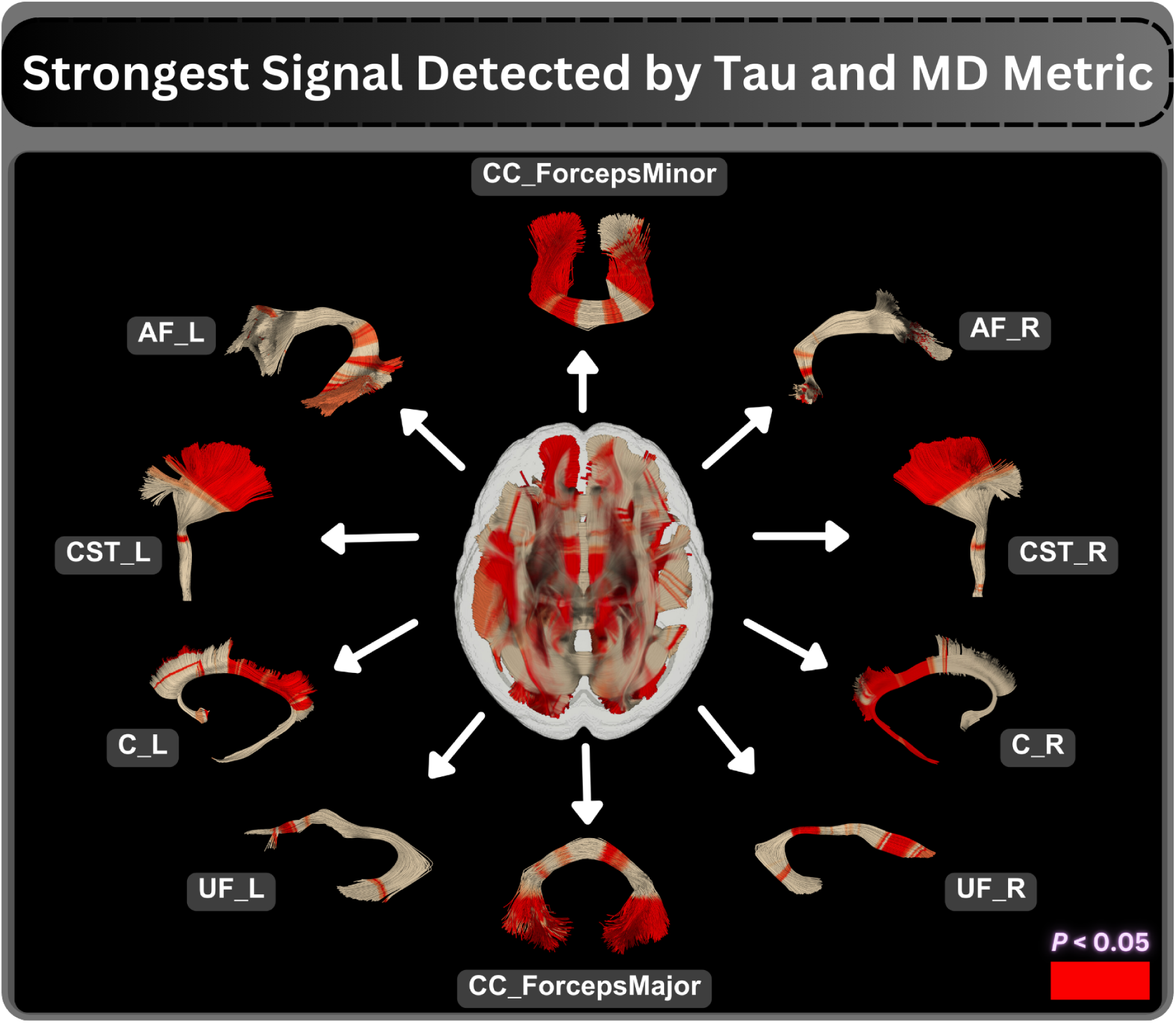
Visualization of localized effects of Tau on MD metrics detected by BUAN. Each tract is colored based on *p*-values, with red indicating p*<*0.05. The combination of Tau and MD detects the strongest association signal among amyloid, tau, ApoE, and the four DTI metrics.

### 3.3. APOE ɛ4 genotype

We ran BUAN to assess the impact of APOE *ɛ*4 - the major common risk gene for late-onset Alzheimer’s disease - on 38 major white matter tracts, based on data from 358 non *ɛ*4 carriers *ɛ*2*ɛ*3/*ɛ*3*ɛ*3/*ɛ*2*ɛ*2 (CN: 224, MCI: 99, Dementia: 35) (168M, 190F) and 203 participants with at least one *ɛ*4 gene; *ɛ*2*ɛ*4/*ɛ*3*ɛ*4/ *ɛ*4*ɛ*4 carriers (CN: 136, MCI: 54, and Dementia: 13) (90M, 108F) participants from the ADNI3 dataset.

Significant effects were detected in the corticospinal tract left (FA), frontopontine tract left (FA), inferior longitudinal fasciculus right (MD), and middle longitudinal fasciculus right (AxD). MD, RD, and AD decrease. FA slightly increases. Fig. 6 highlights the significant tracts, with darker orange indicating areas of significant effects. The color gradient represents *p*values projected onto the 3D tracts, ranging from 1 to 0. Lighter colors indicate *p*values closer to 1 (no significant effects found), while darker colors indicate *p*values less than 0.05.

**Fig. 6.**
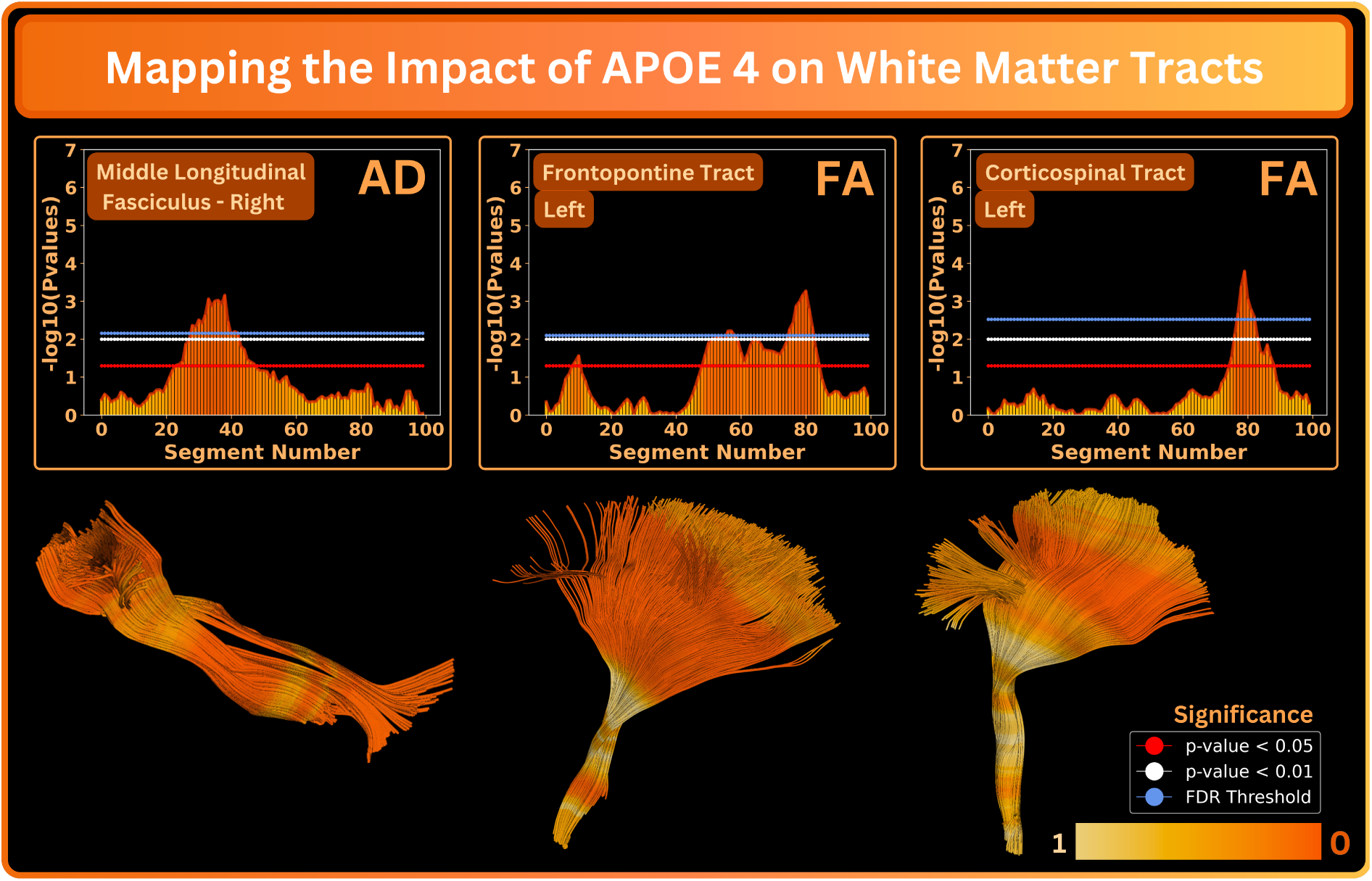
BUAN results for white matter tract alteration in the subgroup of participants carrying at least one *ɛ*4 gene. The first and third rows show *p*value plots for each tract, where the *x*-axis represents the segment number along the tract and the *y*-axis shows a negative logarithm of *p*values. The blue horizontal line in the plots represents the FDR-corrected threshold. Segments that pass the FDR-corrected threshold are considered significant. The second and fourth rows visualize *p*values mapped onto the 3D tracts. Dark orange colors imply lower *p*values and greater strength of association.

### 3.4. APOE ɛ3 vs. APOE ɛ4

We ran BUAN to assess the impact of *ɛ*4 on 38 major white matter tracts using 310 *ɛ*3*ɛ*3 (CN: 191, MCI: 85, Dementia: 34) (140M, 170F) and 192 *ɛ*3*ɛ*4/*ɛ*4*ɛ*4 (CN:129, MCI:50, and Dementia:13) (88M, 104F) subjects from ADNI3 dataset.

Significant findings included frontopontine Tract left (FA), inferior Longitudinal Fasciculus right (AxD, MD), and Middle Longitudinal Fasciculus right (AxD), and spinothalamic tract left (MD), and right (AxD). MD decreases, AD changes vary along the length of the tract, with a slight increase in FA. Tract visualizations shown in Fig. 7 highlight the significant tracts, with darker pink indicating areas of significant effects. The color gradient represents *p*values projected onto the 3D tracts, ranging from 1 to 0. Lighter colors indicate *p*values closer to 1 (no significant effects detected), while darker colors indicate *p*values less than 0.05.

**Fig. 7.**
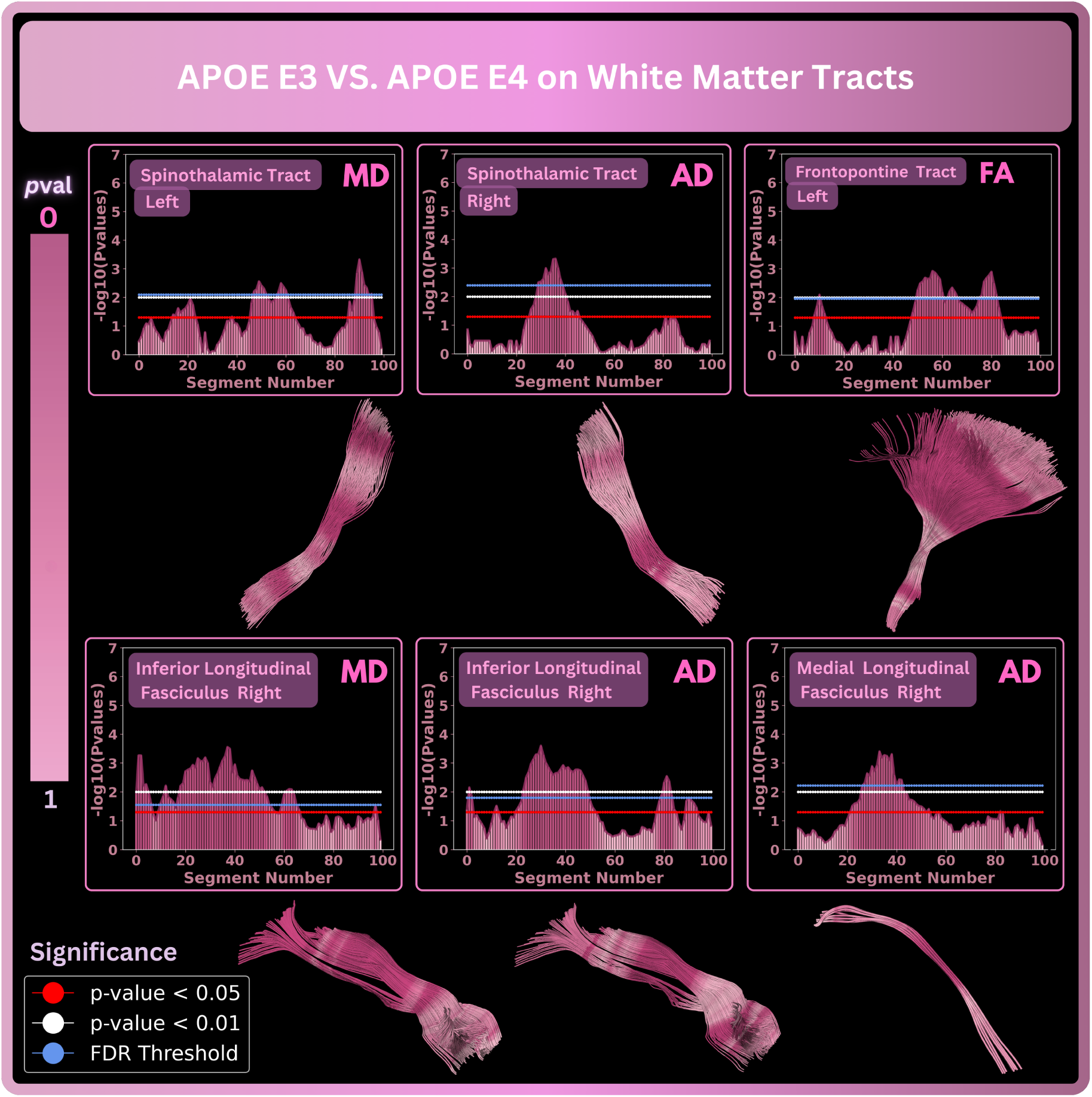
BUAN results for group differences between *ɛ*3*ɛ*3 neutral gene and subjects with either *ɛ*3*ɛ*4 or *ɛ*4*ɛ*4 gene in white matter tracts. The first and third row shows *p*value plots for each tract, where the x-axis represents the segment number along the tract and the y-axis shows a negative logarithm of *p*values. The blue horizontal line in the plots represents the FDR corrected threshold. Segments that pass the FDR corrected threshold are considered significant. The second and fourth rows visualize *p*values mapped onto the 3D tracts. Where dark pink color implies lower *p*values and more significance.

### 3.5. APOE ɛ3 vs. APOE ɛ2

We ran BUAN to assess the impact of the APOE *ɛ*2 genotype (which is protective against Alzheimer’s disease) on 38 major white matter tracts using 310 *ɛ*3*ɛ*3 (CN: 191, MCI: 85, and Dementia: 34) (140M, 170F) and 48 *ɛ*3*ɛ*2/*ɛ*2*ɛ*2 48, (CN: 33, MCI: 14, and Dementia: 1) (28M, 20F) participants in the ADNI3 dataset.

We found the following tracts and measures to be significant: Middle Longitudinal fasciculus right (AxD), spinothalamic tract right (FA, AxD), and uncinate fasciculus right (AxD). FA increases, MD and RD decrease, and AxD changes vary along the length of the tracts. Tract visualizations shown in Fig. 8 highlight the significant tracts, with darker green indicating areas of significant effects. The color gradient represents *p*values projected onto the 3D tracts, ranging from 1 to 0. Lighter colors indicate *p*values closer to 1 (no significant effects found), while darker colors indicate *p*values less than 0.05.

**Fig. 8.**
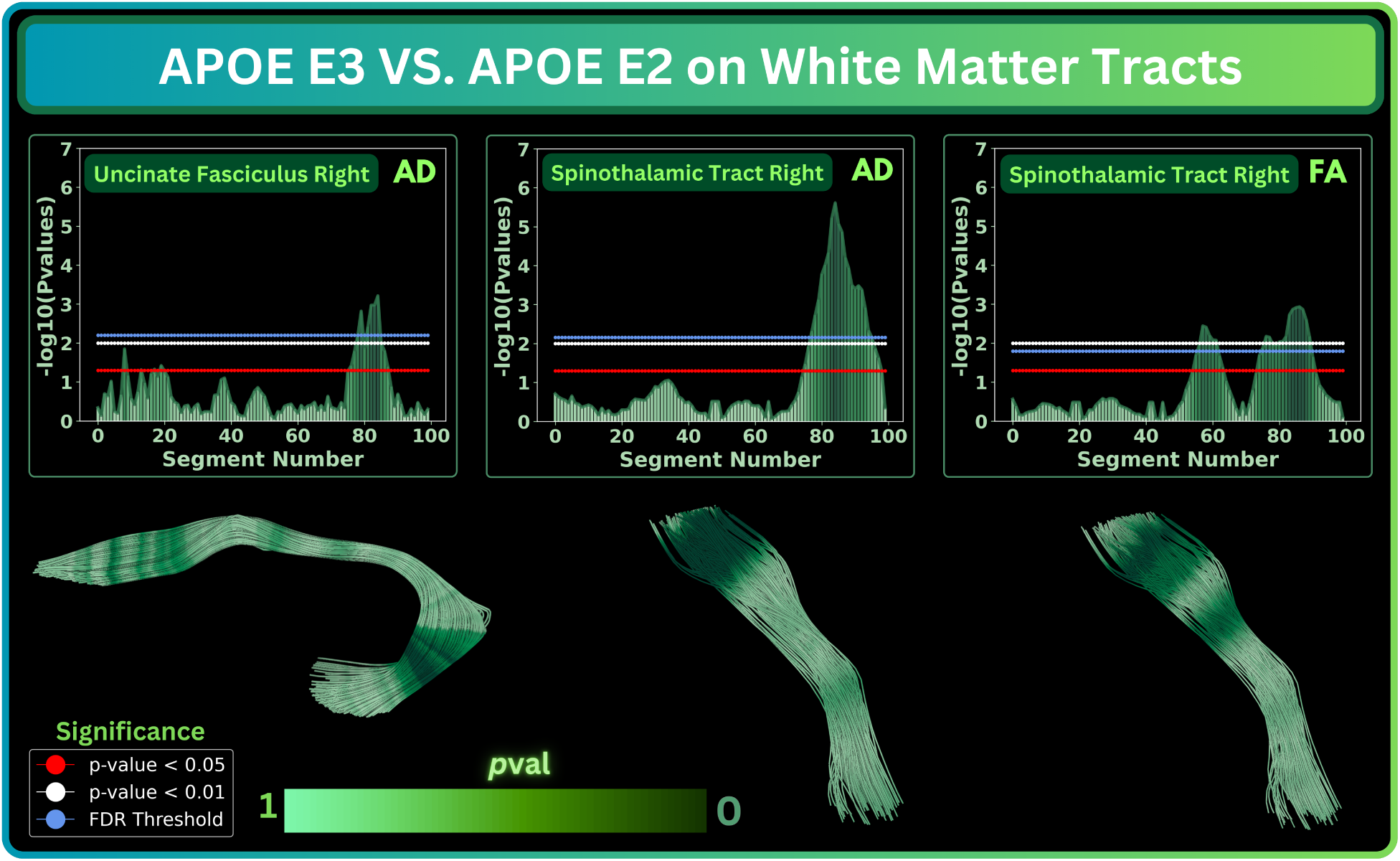
BUAN results for group differences between the *ɛ*3*ɛ*3 (neutral) gene and participants with either *ɛ*3*ɛ*2 or *ɛ*2*ɛ*2 gene in white matter tracts. The first and third rows show *p*value plots for each tract, where the *x*-axis represents the segment number along the tract and the *y*-axis shows a negative logarithm of *p*values. The blue horizontal line in the plots represents the FDR-corrected threshold. Segments that pass the FDR-corrected threshold are considered significant. The second and fourth rows visualize *p*values mapped onto the 3D tracts. Where dark green color implies lower *p*values and greater significance.

## 4. Discussion

Our study employed the advanced tractometry method, BUAN (Bundle Analytics), to investigate the effects of amyloid, tau, APOE *ɛ*4, and APOE *ɛ*2 on the microstructure of white matter tracts in the brain. The results underscore the significant role of tau and amyloid as biomarkers for Alzheimer’s disease (AD), revealing their profound impact on white matter integrity. Tau and amyloid deposition are associated with marked changes in MD, AD, and RD, with FA being the least sensitive metric. This highlights the critical nature of these biomarkers in the early detection and monitoring of AD progression.

Tau and amyloid significantly alter the microstructural properties of white matter tracts, which are essential for neural communication. APOE *ɛ*4 carriers showed microstructural changes consistent with poorer white matter integrity, compared to those with the *ɛ*3/*ɛ*3 genotype, in line with the heightened genetic risk for AD associated with APOE *ɛ*4. These alterations are likely due to the reduced efficiency of amyloid clearance and increased inflammation observed in *ɛ*4 carriers. Conversely, fewer white matter bundles were affected by APOE *ɛ*2, perhaps in line with its protective role against AD-related white matter degeneration.^11,12^ The findings also revealed that MD is the most affected metric, followed by AxD and RD, whereas FA is the least sensitive. This is consistent with prior literature studying the association of DTI metrics with dementia.^65,66^ This differential sensitivity of diffusion metrics highlights the importance of selecting appropriate imaging markers for assessing white matter integrity in AD. MD, in particular, may serve as a more reliable indicator of microstructural damage in the context of AD pathology.

Our results underscore the significant role of the key Alzheimer’s disease biomarkers in altering the microstructure of key neural pathways, with profound implications for understanding the progression and potential intervention points for AD. Some key tracts - the cingulum bundles and components of the corpus callosum - showed significant alterations in MD and RD in the presence of both amyloid and tau. The increased MD and RD indicate water molecules diffusing more freely in brain tissue - a sign of tissue degeneration and loss of cellular integrity typical in Alzheimer’s disease. This diffusion behavior reflects the structural breakdown of neural pathways, which is critical in the progression of Alzheimer’s disease.

Effects of amyloid and tau on crucial neural pathways, such as the cingulum and corpus callosum, may lead to impaired cognitive function and poorer interhemispheric communication. The cingulum bundle is critical for cognitive and emotional processing, and disruption of its microstructure can lead to impaired connectivity between the frontal lobe and other brain regions, contributing to the cognitive decline in AD patients.^67,68^ The corpus callosum (CC), the largest white matter structure in the brain, facilitates inter-hemispheric communication. The impact on the corpus callosum (including forces major, forceps minor, and the middle sector of CC) is of great interest, as it plays a crucial role in coordinating cognitive and motor functions across the two brain hemispheres.^69^ Additional tracts, such as the extreme capsule (EMC) and frontopontine tract (FPT), inferior longitudinal fasciculus (ILF), middle longitudinal fasciculus (ML), optic radiation (OR), and spinothalamic tract (STT), also exhibited significant changes in diffusivity metrics in relation to amyloid and tau. The EMC tract is located in the temporal lobe and is involved in auditory and language processing.^70^ Damage to this tract can affect communication and language abilities, common areas of decline in AD patients. The FPT connects the frontal cortex to the pons,^19^ and its impairment can lead to issues with motor control and executive functions, which are often observed in AD. The ILF connects the temporal and occipital lobes, playing a role in visual processing and memory.^71^ Disruptions in the ILF can contribute to visual memory deficits. The MLF is involved in language, semantic memory, and the integration of auditory and visual information.^72^ Its impairment may contribute to semantic and memory deficits. OR carries visual information from the thalamus to the visual cortex, impairment in this tract can affect visual processing,^73^ which is vital for spatial orientation and navigation. STT is critical for pain and temperature sensation.^74^ While not typically associated with Alzheimer’s core symptoms, its impairment could affect sensory processing. These findings suggest that the effects of AD are not confined to a single functional domain but instead disrupt multiple neural pathways, leading to the diverse clinical manifestations of the disease.

Moreover, this study highlights the limitations of earlier methods such as tract-based spatial statistics (TBSS),^29^ which, despite identifying significant associations between amyloid positivity, clinical dementia severity, and specific brain regions[citations], suffers from limited resolution due to predefined atlas regions. The BUAN method overcomes these limitations by offering a finer-scale mapping of microstructural changes along the length of white matter tracts, providing a more detailed and accurate assessment of disease-related alterations. The pronounced effects detected in specific bundles reveal the vulnerability of these white matter fiber pathways to Alzheimer’s disease pathology, highlighting their potential as biomarkers for early detection and monitoring of disease progression. Understanding how these tracts are compromised allows for more targeted research into therapeutic interventions to preserve or restore their integrity.

Future work will integrate microstructural measures derived from sophisticated modeling techniques, such as diffusion kurtosis imaging (DKI),^75^ or neurite orientation dispersion and density imaging (NODDI)^76^ into BUAN. Notably, NODDI shows superior ability in capturing the microstructural properties of brain tissue, outperforming traditional DTI metrics.^77^

## 5. Conclusion

In this study, we employ our advanced tractometry method, BUAN (Bundle Analytics), to evaluate the impact of amyloid, tau, APOE *ɛ*4, and APOE *ɛ*2 on the microstructural properties of white matter tracts in the brain. Among these factors, we find that microstructural alterations in white matter tracts are most significantly associated with tau and amyloid – the two prominent biomarkers of Alzheimer’s disease. Fewer bundles are affected by APOE *ɛ*2, and comparing APOE *ɛ*4 with APOE *ɛ*3/*ɛ*3 reveals stronger microstructural alterations than comparing APOE *ɛ*4 with *ɛ*2 and *ɛ*3 variants combined.

## 6. Acknowledgement

This research was supported by the NIH (National Institutes of Health) under the AI4AD project grant U01 AG068057, grant numbers P41 EB015922, and RF1 AG057892. We would like to acknowledge the National Institute of Biomedical Imaging and Bioengineering under award numbers R01 EB027585 and R01 EB017230.

## Notes

### Competing Interest Statement

The authors have declared no competing interest.

### Summary of Updates

This version of the manuscript has been revised to update add ADNI {and for the Alzheimer's Disease Neuroimaging Initiative*) consortium as co-author

